# Natural Compound–Mediated SUMOylation Rewires Host Immunity Against *Klebsiella pneumoniae*

**DOI:** 10.64898/2026.09.12.751131

**Authors:** Sandhya Padmakumar, Abhinand Kuniyil, Abhijith Pulimoottil Jaykumar, Athira Gopakumar, Mydhily Soorej, Pradeesh Babu, Bipin G. Nair, Aravind Madhavan, Geetha B. Kumar

**Affiliations:** School of Biotechnology, Amrita University, Amrita Vishwa Vidyapeetham, Clappana P.O., Kollam, Kerala, 690525, India

**Keywords:** Multidrug-resistant hypervirulent *K. pneumoniae*, SUMOylation, post-translational modification, Clove bud oil (CBO), immunomodulatory effect

## Abstract

*Klebsiella pneumoniae* is a major cause of multidrug-resistant and hypervirulent infections worldwide, highlighting the urgent need for host-directed therapeutic strategies that complement conventional antibiotics. Emerging evidence suggests that *K. pneumoniae* promotes intracellular persistence by suppressing host SUMOylation, a reversible post-translational modification that regulates protein stability, cellular signaling, and innate immune response. This decrease in SUMOylation perturbs NF-κB signaling, leading to dysregulation of inflammation that promotes bacterial survival, rather than effective immune clearance. Clove Bud Oil (CBO), a natural product derived from *Syzygium aromaticum*, has previously been shown to possess anti-virulence as well as immunomodulatory properties. In this regard, we investigated the capability of CBO to restore host SUMOylation and enhance innate immune response during infection with classical as well as hypervirulent *K. pneumoniae*. In macrophages, classical and hypervirulent *K. pneumoniae* impaired global host SUMOylation, disrupting IKKβ–RelA/NF-κB signalling and uncoupling inflammatory regulation from effective antimicrobial defense to facilitate intracellular persistence. CBO restored SUMOylation-dependent immune homeostasis by attenuating sustained IKKβ phosphorylation and degradation, thereby constraining aberrant RelA nuclear translocation, rebalancing pro-inflammatory cytokine responses, and restricting intracellular bacterial survival. These findings demonstrate that restoration of host SUMOylation acts as a key driver of the immunomodulatory activity of CBO, leading to coordinated innate immune responses and enhanced intracellular bacterial clearance. Collectively, our study identifies host SUMOylation as a critical pathway targeted during *K. pneumoniae* infection and highlights restoration of SUMOylation-mediated immune homeostasis as a promising host-directed therapeutic strategy against multidrug-resistant and hypervirulent *K. pneumoniae*.

**Author Summary:** Antibiotic-resistant *Klebsiella pneumoniae* has become a major global health threat, making it increasingly important to identify alternate strategies to treat infections that do not rely solely on antibiotics. In this study, we investigated how *Klebsiella pneumoniae* interferes with host immune defenses. Our findings demonstrate that the pathogen disrupts SUMOylation, a key cellular process involved in coordinating balanced immune responses. Impaired SUMOylation dysregulates immune signaling, thereby promoting bacterial survival within host cells. We further examined whether clove bud oil (CBO), a natural product with established biological activity, could restore SUMOylation-mediated immune regulation. Our results show that clove bud oil re-established host SUMOylation, normalized immune signaling, reduced excessive inflammatory responses, and improved the ability of immune cells to eliminate both classical and hypervirulent strains of *K. pneumoniae*. These findings suggest that the beneficial effects of clove bud oil arise not only from its antimicrobial properties but also from its ability to modulate the host immune response by restoring normal cellular regulation. Our study identifies host SUMOylation as a promising therapeutic target and provides evidence that strengthening the body’s own immune defenses may offer an alternative strategy to combat multidrug-resistant bacterial infections.

## 1. Introduction

The emergence of Anti-Microbial Resistance (AMR) is escalating as a global health crisis that severely compromises the effectiveness of existing antibiotic therapies. Among bacterial infections, Gram-negative pathogens have emerged as leading causes of multidrug-resistant (MDR) infections, particularly in respiratory diseases, including community-acquired pneumonia, hospital-acquired pneumonia, and ventilator-associated pneumonia [1]. The growing prevalence of hypervalent eclipse pneumonia, HVP strain, further intensifies this threat, as these lineages exhibit enhanced immune evasion, intracellular persistence, and tissue dissemination, even in immunocompetent individuals [2]. The coverage of multidrug resistance and increased virulence in cellular systems urgently requires alternative therapeutic approaches that go beyond direct bacterial killing to strengthen host immune defense [3].

Host-directed therapies (HDTs) have gained increasing attention as a complementary strategy to conventional antibiotics, as they aim to restore immune homeostasis by minimizing selective pressure for resistance [4]. Among the various therapeutic strategies, natural products, such as essential oils, have emerged as promising candidates due to their diverse biological activities, including anti-virulent, anti-inflammatory, and immunomodulatory properties. Clove bud oil (CBO), derived from Syzygi*um aromaticum*, has a long history of medicinal use and is known for its antimicrobial and immune regulatory properties [5]. Previous work from our laboratory and others has demonstrated that CBO can attenuate bacterial virulence [6] and moderate macrophage immune responses, highlighting its potential as a host-directed modulator against bacterial infections.

Post-translational modification systems constitute critical regulatory nodes during host-pathogen interactions and are frequently targeted by bacterial pathogens to subvert immune responses [7]. SUMOylation, a reversible modification involving the conjugation of Small Ubiquitin-like Modifier (SUMO) proteins to target substrates, regulates diverse cellular processes, including protein stability, subcellular localization, transcriptional activity, and signal transduction [8]. In immune cells, SUMOylation acts as a critical regulator of protein stability, localization, and function by orchestrating immune signalling pathways, thereby influencing the host’s antimicrobial defense mechanisms [9]. Notably, *Klebsiella pneumoniae* infection has been shown to induce a global reduction in host immunity, thereby impairing effective inflammatory responses and promoting intracellular survival of these pathogens within macrophages [10].

Among the several signalling pathways influenced by SUMOylation, NF-κB plays a central role in coordinating innate immune responses during infection. Core components of this pathway, including IκB kinase β (IKKβ) and the NF-κB subunit RelA (p65), are subject to dependent phosphorylation that modifies their association status and nuclear localization [11]. Disruption of SUMOylation during *K. pneumoniae* infection destabilizes IKKβ, leading to NF-κB activation and deregulated cytokine production that favors bacterial persistence [12]. The study aims to determine whether CBO restores host SUMOylation during *Klebsiella pneumoniae* infection and modulates the crosstalk between SUMOylation and NF-κB signalling to re-establish immune homeostasis. By elucidating the molecular mechanisms, our work focuses on SUMOylation as a key host regulatory pathway during respiratory infection. It proposes clove oil as a promising host-directed therapeutic agent to enhance host defense and mitigate infection-induced inflammation.

## 2. Results

### 2.1. Growth kinetic analysis identifies Clove Bud Oil (CBO) as the most effective inhibitor of *Klebsiella pneumoniae* growth

Growth kinetic analysis was performed to evaluate the antibacterial activity of CBO against the classical *Klebsiella pneumoniae* strain ACC33495 and a hypervirulent *K. pneumoniae* strain. CBO exhibited a concentration-dependent inhibitory effect on bacterial growth over the 12 h incubation period. Increasing CBO concentrations progressively suppressed bacterial proliferation, with 0.25% CBO showing the greatest inhibition compared with the untreated control. Similar growth inhibition was observed in both the classical and hypervirulent strains, indicating the broad-spectrum antibacterial activity of CBO **(Figure S2)** .

### 2.2. Selection of a non-cytotoxic concentration of CBO for macrophage infection studies

To identify a physiologically relevant concentration of CBO suitable for host-directed studies, THP-1 monocytes were differentiated into macrophages and exposed to increasing concentrations of CBO for 24 h and 48 h, followed by MTT analysis. It was observed that THP-1 cells were well tolerated at concentrations of 0.01–0.015%, with high cell viability maintained, whereas concentrations ≥0.02% resulted in a marked, dose-dependent reduction in viability **(Figure 1A)**. To further validate the host safety of CBO beyond THP-1 cells, primary human PBMCs were exposed to increasing concentrations of CBO and assessed by MTT assay. PBMC viability also remained high at 0.01–0.015% CBO, confirming these concentrations have minimal cytotoxicity and can be used for functional immune assays. In contrast, higher concentrations resulted in a dose-dependent reduction in PBMC viability. These results confirm that 0.015% CBO is well tolerated in both immortalized macrophages and primary immune cells, strengthening its relevance as a host-directed immunomodulatory agent **(Figure 1C)**.

**Figure 1.**
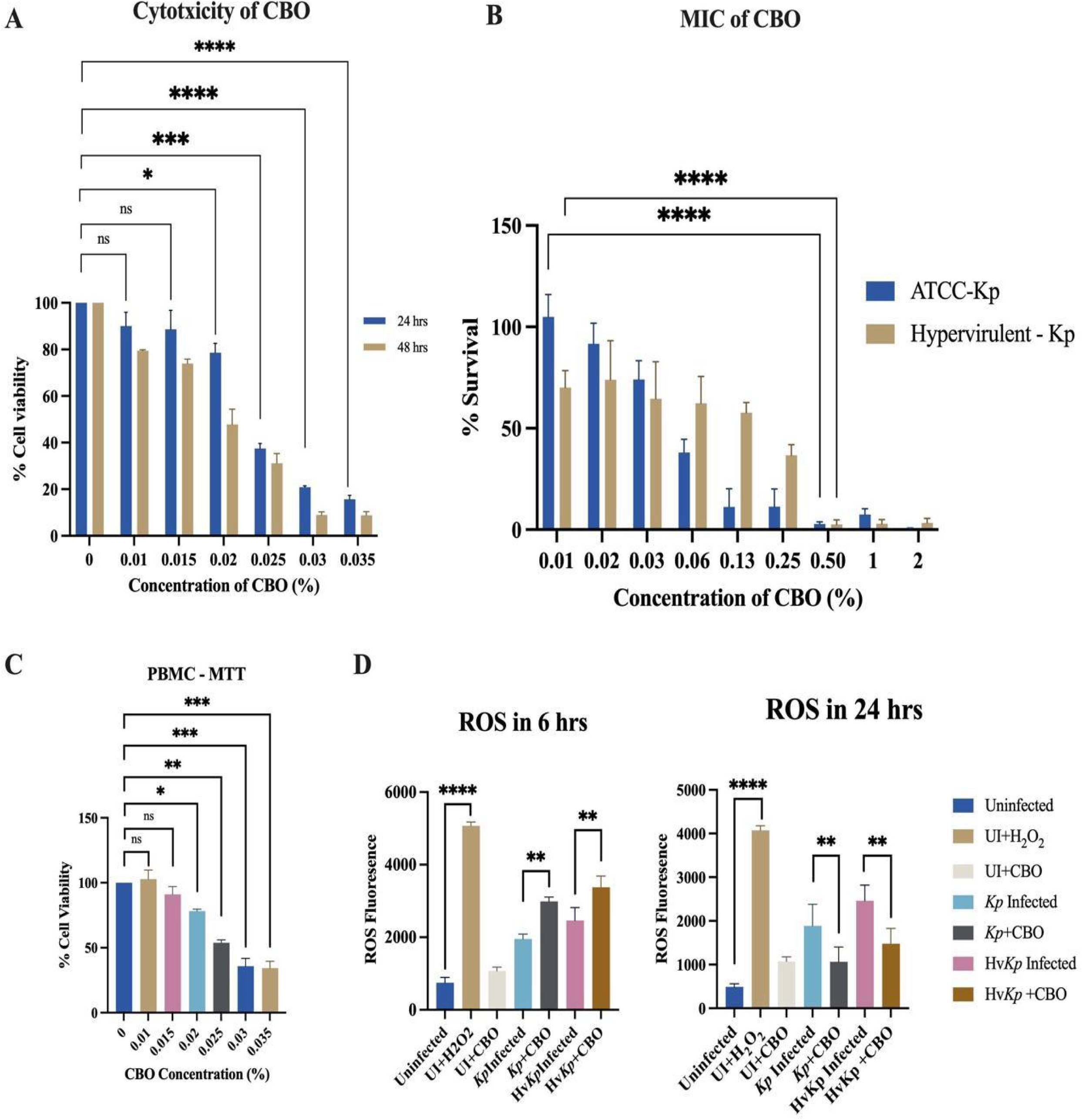
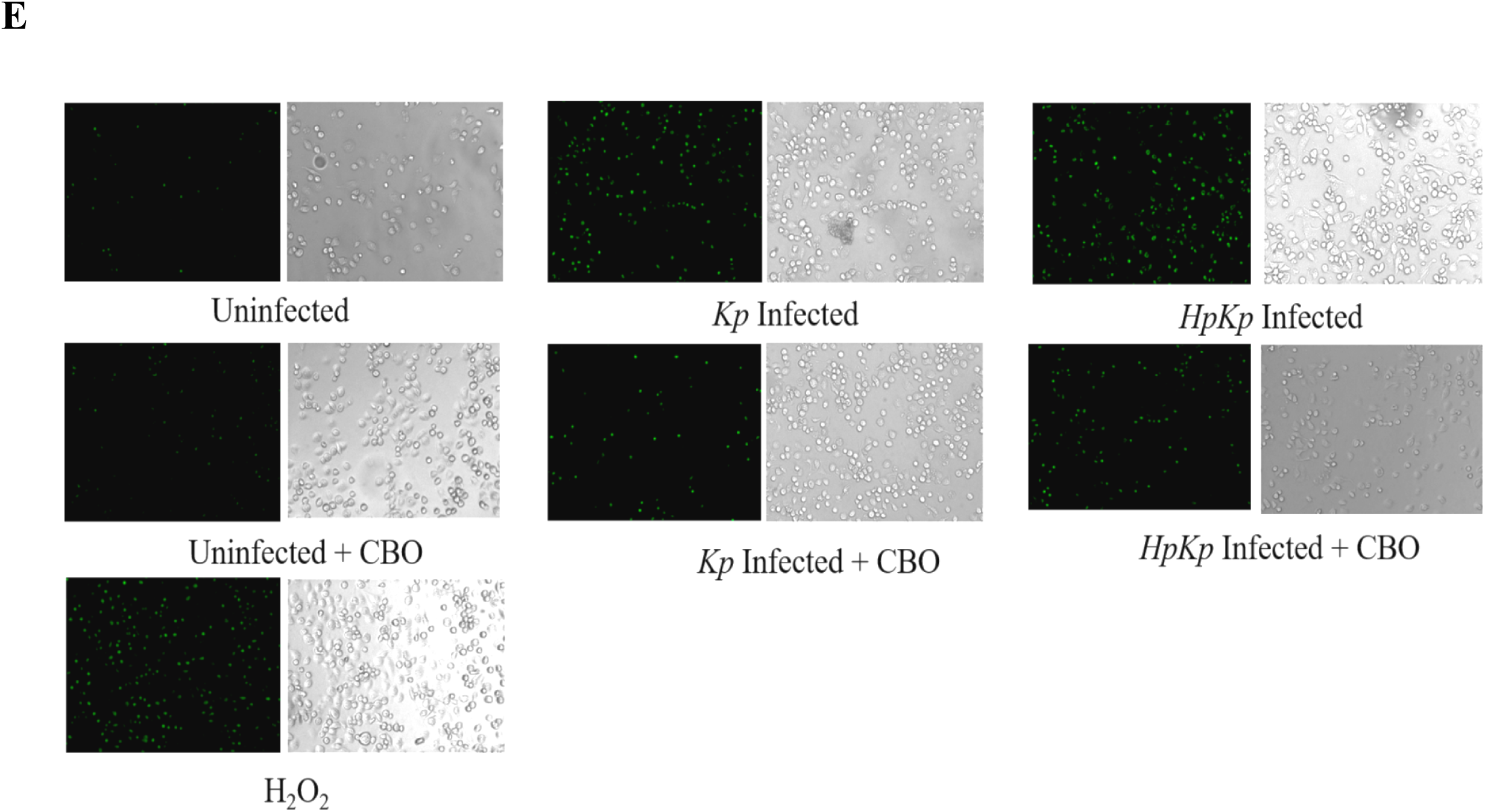
Cytotoxicity, antibacterial activity, and immunomodulatory effects of Clove Bud Oil (CBO). (A) Cytotoxicity of CBO in THP-1-derived macrophages following 24 h and 48 h treatment, assessed by MTT assay. (B) Minimum inhibitory concentration (MIC) analysis of CBO against *Klebsiella pneumoniae* ATCC 33495 and hypervirulent *K. pneumoniae* (hvKp). (C) Cytotoxicity of CBO in primary human PBMCs determined by MTT assay. (D) Intracellular ROS production was measured by DCFH-DA assay at 6 h and 24 h post-infection, with H₂O₂ as the positive control. (E) Representative fluorescence images showing intracellular ROS levels under the indicated conditions. Data are presented as mean ± SD from independent experiments. Statistical significance was determined by one-way ANOVA; *P* < 0.05 was considered significant.

### 2.3. CBO exhibits concentration-dependent growth inhibition of classical and hypervirulent *K. pneumoniae*

The minimum inhibitory concentration (MIC) of CBO was determined against classical *Klebsiella pneumoniae* (ATCC) and hypervirulent *K. pneumoniae* (hvKp). CBO exhibited concentration-dependent antibacterial activity against both strains, with the hypervirulent strain showing slightly greater tolerance at intermediate concentrations. The MIC90 for classical and hypervirulent *K. pneumoniae* was 0.1% CBO **(Figure 1B)**. Based on the cytotoxicity and MIC analyses, a sub-MIC concentration of 0.015% CBO was selected for all subsequent host–pathogen experiments.

### 2.4. CBO enhances early intracellular ROS generation during *K. pneumoniae* infection

Intracellular reactive oxygen species (ROS) production was assessed using the DCFH-DA assay to evaluate the contribution of oxidative antimicrobial responses during infection and treatment. Initial fluorescence microscopy images acquired at 24 h post-infection revealed relatively low ROS levels in CBO-treated infected macrophages compared with earlier expectations. As ROS generation represents an early antimicrobial defense mechanism that typically occurs during active pathogen clearance, a shorter time point was subsequently investigated to better capture the oxidative burst associated with bacterial killing.

Analysis at 6 h post-treatment demonstrated a marked increase in intracellular ROS production in CBO-treated infected macrophages compared with infected controls for both classical and hypervirulent *K. pneumoniae* infections. In contrast, infection alone failed to induce a comparable oxidative response. The increased ROS generation observed at the early time point suggests that CBO enhances macrophage antimicrobial activity during the active host–pathogen interaction phase. We understand the reduced ROS signal observed at 24 h likely reflects a later stage of infection following bacterial clearance and resolution of the oxidative burst **(Figure 1D)**. Representative DCFH-DA fluorescence images were acquired at 24 h post-infection, whereas quantitative ROS measurements were used to evaluate early oxidative responses. Collectively, these findings indicate that CBO promotes an early ROS-mediated antimicrobial response that may contribute to enhanced intracellular bacterial clearance **(Figure 1E)**.

### 2.5. Effect of Clove Bud Oil on Phagocytosis and Intracellular Survival of Klebsiella pneumoniae

#### 2.5.1. Effect of CBO on Phagocytosis

To determine whether CBO enhances host cell uptake of *K. pneumoniae*, phagocytosis assays were performed using both ATCC and hypervirulent *K. pneumoniae* (hvKp) strains in THP1 cells and Human PBMCs **(Figure 2A, 2D)**. Infection alone resulted in intracellular bacterial counts of approximately 0.8 × 10⁶ CFU/ml for both strains. In contrast, CBO-treated cells showed significantly higher intracellular bacterial recovery, reaching approximately 2.2 × 10⁶ CFU/ml for the ATCC strain and 2.7 × 10⁶ CFU/ml for the hvKp strain. These results indicate that CBO markedly enhances bacterial uptake by host cells, suggesting improved phagocytic activity. The enhancement was observed for both classical and hypervirulent strains, demonstrating that the effect of CBO is not strain-specific. Increased bacterial internalization suggests that CBO may stimulate host cell functions involved in pathogen recognition and uptake.

**Figure 2.**
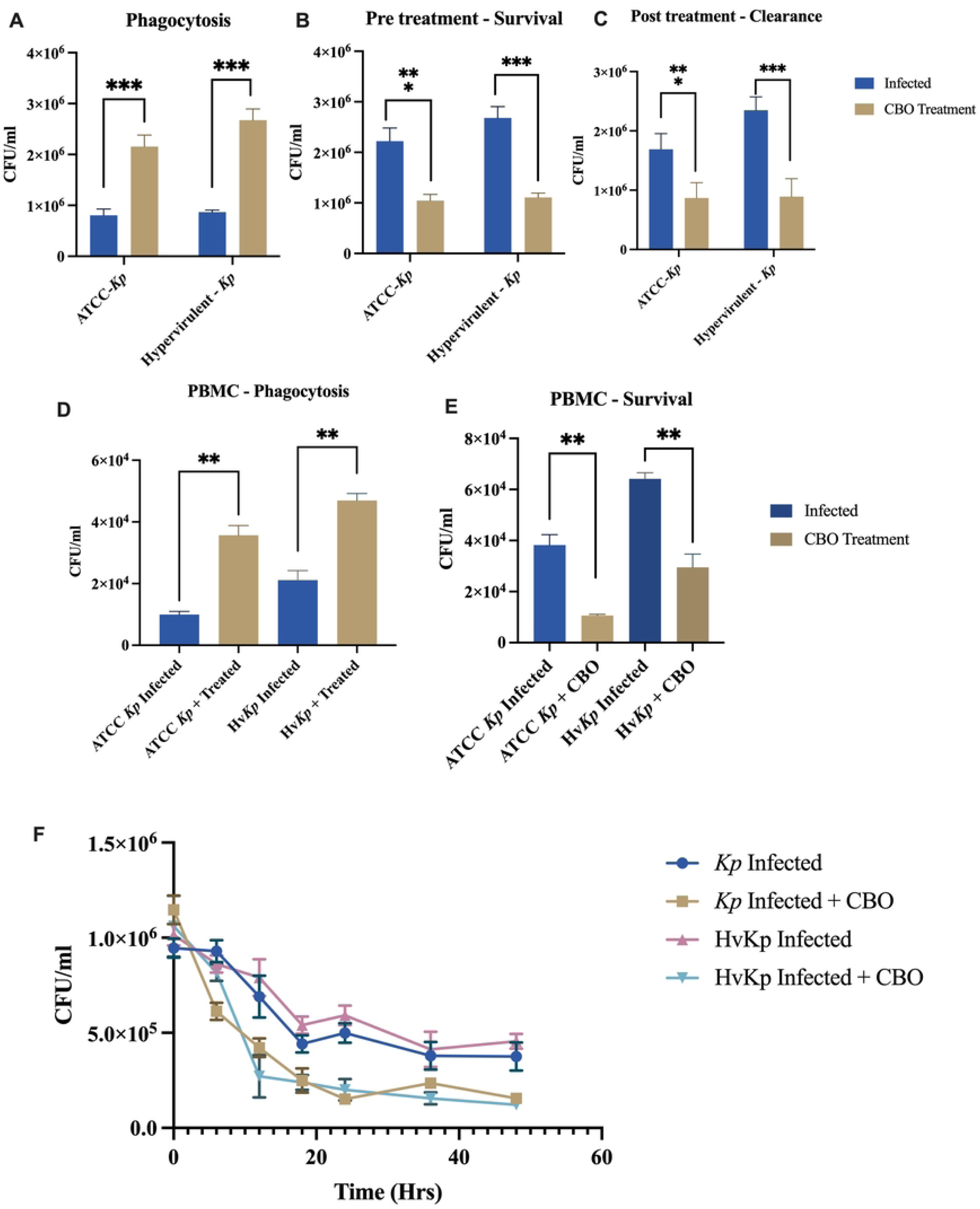
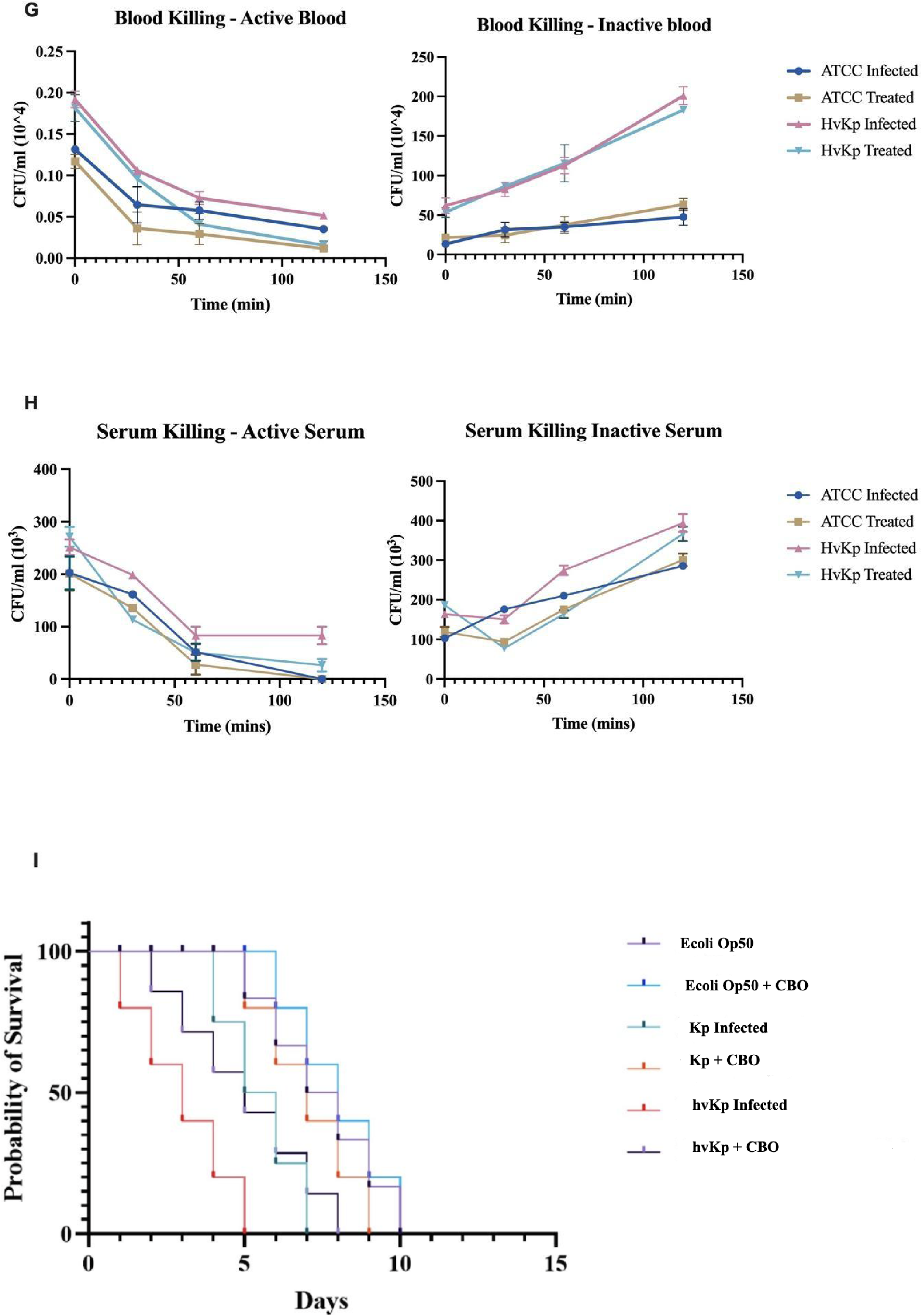
Clove bud oil enhances host-mediated clearance of Klebsiella pneumoniae across multiple infection models: (A) Phagocytosis assay in THP-1 macrophages. (B) Pre-treatment intracellular survival assay. (C) Post-treatment intracellular clearance assay. (D) PBMC phagocytosis assay. (E) PBMC intracellular survival assay. (F) Time-course intracellular killing assay showing accelerated bacterial clearance following CBO treatment. (G) Whole-blood killing assays performed in active and heat-inactivated blood. (H) Serum killing assays performed in active and heat-inactivated serum. (I) Kaplan–Meier survival analysis of *C. elegans* infected with ATCC or hypervirulent *K. pneumoniae* (hvKp). CBO treatment enhanced bacterial uptake, reduced intracellular survival, improved complement-dependent bacterial killing, and prolonged host survival. Data are presented as mean ± SEM. Statistical significance is indicated as P < 0.05, P < 0.01, or P < 0.001.

#### 2.5.2. Effect of CBO Pre-treatment on Intracellular Survival

To evaluate whether prior exposure to CBO could improve host-cell resistance to infection, cells were pretreated with CBO before bacterial challenge **(Figure 2B)**. In the ATCC strain, infected cells exhibited intracellular bacterial counts of approximately 2.2 × 10⁶ CFU/ml, whereas CBO pre-treatment reduced the bacterial burden to nearly 1.0 × 10⁶ CFU/ml. Similarly, hvKp-infected cells showed approximately 2.7 × 10⁶ CFU/ml, which decreased to about 1.1 × 10⁶ CFU/ml following CBO pre-treatment. The substantial reduction in intracellular bacterial burden indicates that pre-exposure to CBO enhances host cells’ ability to control subsequent infection. These findings suggest a prophylactic or immune-priming effect of CBO that prepares host cells to respond more efficiently to bacterial invasion.

#### 2.5.3. Effect of CBO on Intracellular Bacterial Clearance

The ability of CBO to promote clearance of already established intracellular infection was assessed by treating infected cells after bacterial uptake **(Figure 2C)**. In ATCC-infected cells, bacterial counts decreased from approximately 1.7 × 10⁶ CFU/ml to 0.9 × 10⁶ CFU/ml following treatment. Similarly, hvKp-infected cells showed a reduction from approximately 2.3 × 10⁶ CFU/ml to 0.9 × 10⁶ CFU/ml after CBO exposure. These findings demonstrate that CBO not only promotes bacterial uptake but also enhances intracellular killing and clearance. The reduction in bacterial burden following treatment suggests activation of host antimicrobial mechanisms that restrict bacterial persistence within infected cells.

### 2.6. Effect of CBO on PBMC-Mediated Phagocytosis and Bacterial Survival

To validate the observations in primary immune cells, phagocytosis and survival assays were performed using PBMCs **(Figures 2D, 2E)**. Like macrophage cells, CBO treatment significantly increased bacterial uptake in PBMCs. ATCC-infected PBMCs showed bacterial counts of approximately 1.0 × 10⁴ CFU/ml, which increased to 3.5 × 10⁴ CFU/ml after CBO treatment. Likewise, hvKp uptake increased from approximately 2.1 × 10⁴ CFU/ml to 4.7 × 10⁴ CFU/ml. Despite the increased uptake, intracellular survival was significantly reduced following CBO treatment. ATCC-infected PBMCs exhibited a decrease from approximately 3.8 × 10⁴ CFU/ml to 1.0 × 10⁴ CFU/ml, whereas hvKp survival decreased from approximately 6.4 × 10⁴ CFU/ml to 3.0 × 10⁴ CFU/ml. These findings indicate that CBO simultaneously enhances phagocytic uptake and intracellular bacterial killing in immune cells, supporting its immunomodulatory potential.

### 2.7. Time-Kill Kinetics of Intracellular Bacterial Clearance

To further examine the kinetics of bacterial elimination, intracellular survival was monitored over 48 hours **(Figure 2F)**. Both ATCC and hvKp strains exhibited a gradual decline in bacterial burden over time; however, CBO-treated groups showed a markedly faster reduction. The most pronounced decrease was observed within the first 12–18 hours following treatment. Throughout the study period, bacterial counts remained consistently lower in the CBO-treated groups compared to untreated infected controls. These results demonstrate that CBO accelerates bacterial clearance and limits intracellular persistence in both classical and hypervirulent *K. pneumoniae* infections. The sustained reduction in bacterial burden over time further supports the role of CBO as an effective host-directed immunomodulatory agent. Therefore, CBO at the selected sub-MIC concentration (0.015%) significantly enhanced phagocytosis, improved intracellular bacterial killing, reduced bacterial survival in both epithelial cells and PBMCs, and accelerated bacterial clearance over time. Collectively, these findings established 0.015% CBO as an ideal concentration for subsequent mechanistic studies, as it enhanced host-mediated bacterial clearance without exerting direct antibacterial or cytotoxic effects, thereby allowing its immunomodulatory effects on host SUMOylation and innate immune signaling to be investigated.

### 2.8. Whole-blood killing assay: complement-dependent control of *K. pneumoniae* and effect of treatment

To evaluate the effect of CBO on host-mediated bacterial clearance, whole blood killing assays were performed using active and heat-inactivated blood. In active blood, bacterial counts decreased progressively over time in both ATCC and hypervirulent *K. pneumoniae* groups, indicating effective complement-mediated killing. This reduction was more pronounced in the CBO-treated groups. In contrast, bacterial counts increased in heat-inactivated blood, confirming the importance of complement activity for bacterial clearance. Although CBO-treated groups showed slightly lower bacterial counts, bacterial survival remained higher than in active blood. These findings suggest that CBO enhances complement-dependent host defense mechanisms against both classical and hypervirulent *K. pneumoniae* strains **(Figure 2G)**.

### 2.9. Serum Killing assay demonstrates complement-dependent bactericidal activity and the effect of Clove Bud Oil (CBO)

The serum killing assay was performed to determine the role of serum complement and the effect of CBO on bacterial survival. In active serum, bacterial counts decreased steadily over time, with a greater reduction observed in CBO-treated groups compared to infected controls. Both ATCC and hypervirulent strains showed enhanced clearance following treatment. In contrast, bacterial counts increased in heat-inactivated serum, indicating loss of complement-mediated bactericidal activity. Under these conditions, CBO produced only minimal effects on bacterial survival. These results demonstrate that complement plays a critical role in serum-mediated killing and suggest that CBO enhances bacterial clearance by supporting complement-dependent immune responses **(Figure 2H)**.

### 2.10. Clove Bud Oil Improves *C. elegans* Survival During *K. pneumoniae* Infection

The protective effect of CBO was further evaluated using a *C. elegans* infection model. Infection with both ATCC and hypervirulent *K. pneumoniae* significantly reduced worm survival, with hvKp causing more rapid mortality. Treatment with CBO improved survival in both infection groups and delayed mortality compared with untreated controls. The survival benefit was particularly evident in the hvKp-infected worms, where CBO extended lifespan despite the higher virulence of the strain. These findings indicate that CBO reduces bacterial pathogenicity and enhances host resistance to infection. However, the enhanced phagocytosis, reduced intracellular bacterial survival, improved complement-mediated killing, and prolonged host survival collectively support the immunomodulatory potential of CBO against both classical and hypervirulent *K. pneumoniae* infections **(Figure 2I)**.

### 2.11. CBO upregulates SUMO pathway gene expression in infected macrophages

To determine whether CBO could modulate SUMO genes and if it could reflect transcriptional control of the conjugation machinery, we quantified the SUMO pathway genes. The observation showed that the infection downregulated SUMO1, UBCA2, the SUMO E1 gene, and UBC9, the SUMO E2 gene-related transcripts, whereas CBO treatment significantly increased their expression in infected cells, restoring or exceeding baseline levels. This coordinated upregulation supports the idea that CBO strengthens SUMOylation capacity at multiple layers **(Figure 3A-3C).**

**Figure 3:**
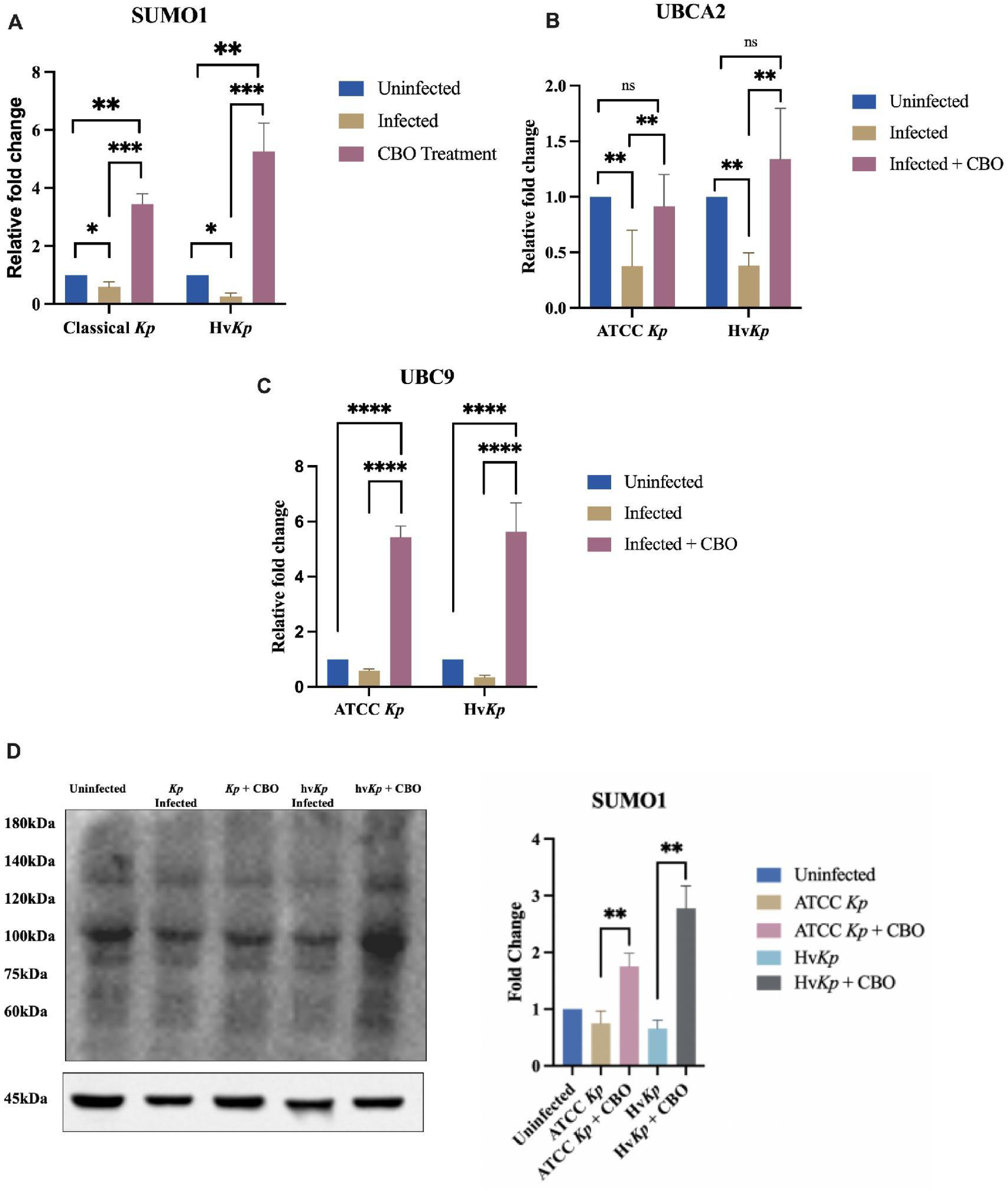
Clove bud oil restores the host SUMOylation machinery during Klebsiella pneumoniae infection: (A–C) Relative mRNA expression of SUMO1 (A), UBA2 (B), and UBC9 (C) in THP-1 macrophages infected with classical *K. pneumoniae* (Kp) or hypervirulent *K. pneumoniae* (hvKp) following CBO treatment. Infection suppressed the expression of SUMOylation-associated genes, whereas CBO significantly restored their expression. (D) Representative immunoblot and densitometric quantification of global SUMO1-conjugated proteins. CBO treatment restored SUMO1 conjugation in both Kp- and hvKp-infected cells, indicating recovery of host SUMOylation. Data are presented as mean ± SEM. Statistical significance is indicated as P < 0.05, P < 0.01, P < 0.001, and P < 0.0001.

### 2.12. CBO restores host SUMOylation during Kp and hvKp infection

Global SUMOylation was markedly reduced following both ATCC and hypervirulent *K. pneumoniae* infection, as evidenced by the diminished intensity of SUMO1-conjugated protein species across a broad molecular weight range, indicating pathogen-mediated disruption of host SUMO signaling. Treatment with CBO significantly restored the abundance of SUMO-conjugated proteins in infected cells, with a more pronounced effect observed in the hvKp-infected group. Densitometric analysis confirmed a significant increase in global SUMO1 conjugation following CBO treatment compared with infected controls. These findings suggest that CBO effectively reverses infection-induced suppression of the SUMOylation machinery, thereby re-establishing host post-translational modification networks that are likely required for optimal regulation of innate immune signaling during bacterial infection **(Figure 3D)**.

### 2.13. Integrated analyses identify NF-κB signalling as a prominent SUMOylation-associated pathway

To investigate signaling pathways associated with the host SUMOylation system, a combination of pathway enrichment, Reactome pathway mapping, protein–protein interaction (PPI), and SUMOylation site prediction analyses was performed. Reactome pathway analysis revealed extensive interactions between SUMOylation-associated proteins and components of immune signaling pathways **(Figure 4A)**. Specifically, SUMO1, SUMO2/3, UBE2I (UBC9), UBA2, and PIAS family proteins were linked to key NF-κB signaling regulators, including RELA, NFKBIA, IKBKB, IKBKG, and IKBKE. Further pathway mapping demonstrated a direct SUMOylation relationship between TOPORS, SUMO1, UBE2I, and IKKβ, suggesting that IKKβ may be regulated through SUMO-dependent mechanisms **(Figure 4B)**. Subcellular interaction analysis further illustrated the spatial association of SUMOylation machinery with NF-κB pathway components within both the cytosolic and nuclear compartments **(Figure 4C)**. Several SUMO-conjugating and SUMO-ligase proteins were found in close proximity to RELA and NF-κB regulatory proteins, supporting a potential role for SUMOylation in modulating NF-κB signaling dynamics. Functional enrichment analysis identified significant overrepresentation of multiple infection- and inflammation-related pathways **(Figure 4D)**. Among the top enriched pathways were NF-κB signaling, Toll-like receptor signaling, NOD-like receptor signaling, C-type lectin receptor signaling, and several bacterial and parasitic infection pathways, including shigellosis, salmonellosis, pertussis, legionellosis, and leishmaniasis. These findings indicate a strong association between SUMOylation networks and innate immune responses. Protein–protein interaction analysis demonstrated extensive connectivity between SUMOylation-associated proteins and inflammatory signaling molecules **(Figure 4E)**. Central network hubs included SUMO1, UBA2, UBE2I, RELA, IKBKB, NFKB1, NFKBIA, and IKBKB, suggesting coordinated regulation between SUMOylation and NF-κB signaling pathways. Network simplification further highlighted RELA and IKKβ as major connecting nodes linking SUMOylation components with inflammatory mediators such as TLR4, MYD88, TNF, IL6, and IL1B **(Figure 4F)**. Given the central position of IKKβ within the interaction network, its potential regulation by SUMOylation was further examined using the GPS-SUMO prediction platform. Multiple putative SUMOylation sites were identified within the IKKβ protein sequence, including residues K147, K310, K480, K531, K628, K704, and K756 **(Figure 4G)**. Several of these sites displayed high prediction scores above the selected threshold, supporting the possibility that IKKβ may undergo SUMO-dependent post-translational modification. Collectively, these integrated analyses identify NF-κB signaling as a prominent pathway associated with SUMOylation and suggest that IKKβ may serve as a critical molecular link between SUMOylation machinery and inflammatory signaling responses.

**Figure 4:**
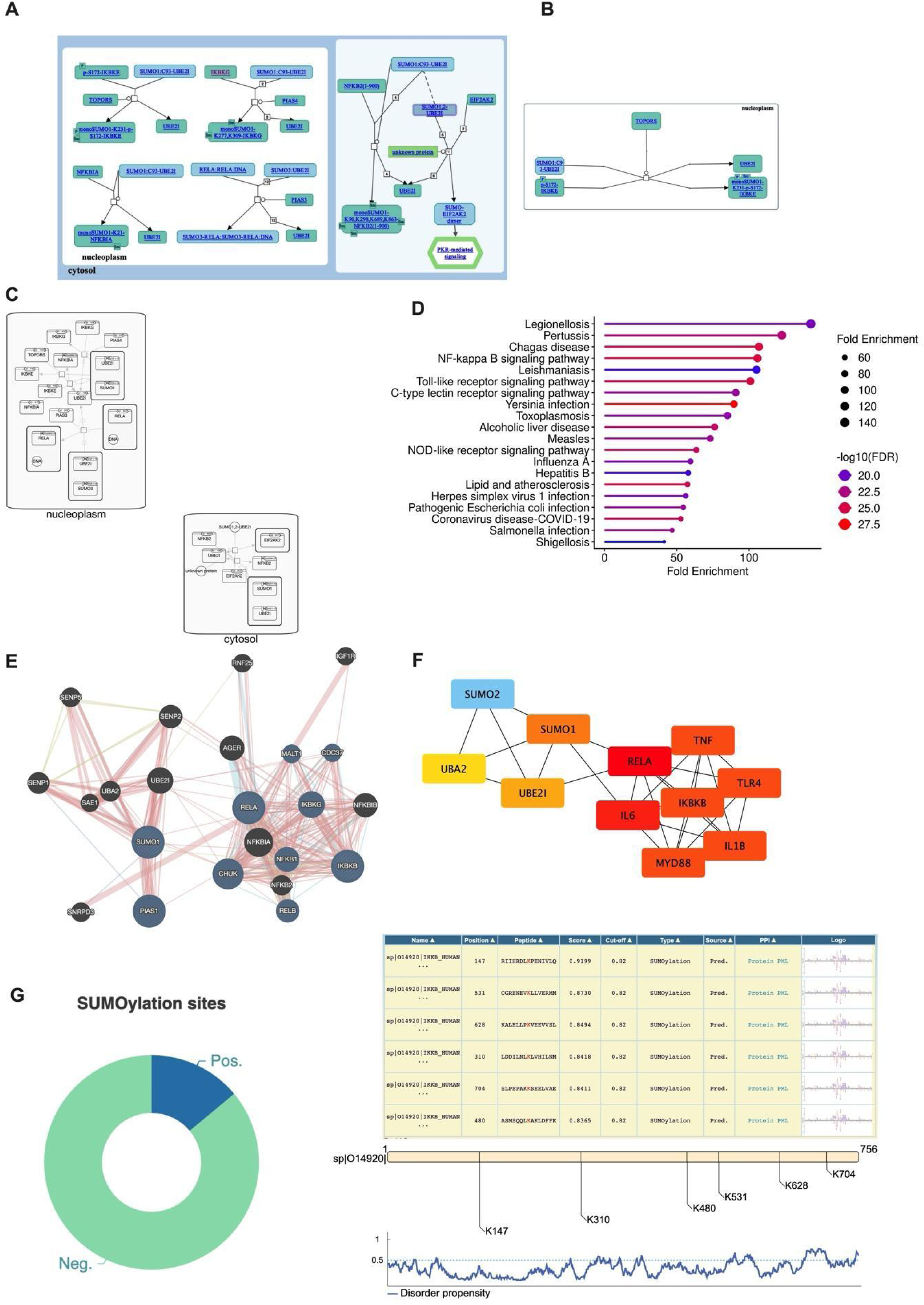
Integrated bioinformatic analyses identify NF-κB signaling and Ikkβ as potential SUMOylation-associated regulatory nodes: (A) Reactome pathway analysis showing SUMOylation-associated immune response pathways and interactions between SUMO machinery components and inflammatory signaling proteins. (B) Reactome interaction map predicting SUMO1-mediated SUMOylation of IKKβ (IKBKB). Subcellular interaction network illustrating the localization of SUMOylation-associated proteins within the cytosolic and nuclear compartments. Functional enrichment analysis of SUMOylation-related genes demonstrating significant enrichment of immune and inflammatory pathways, including NF-κB, Toll-like receptor, and NOD-like receptor signaling pathways. (E) Protein–protein interaction (PPI) network showing extensive connectivity between SUMOylation machinery proteins and NF-κB signaling components. (F) Network analysis highlighting interactions among SUMO pathway proteins (SUMO1, SUMO2, UBA2, and UBE2I) and key inflammatory mediators, including RELA, IKBKB, TLR4, MYD88, TNF, IL6, and IL1B. (G) GPS-SUMO prediction analysis identifying multiple putative SUMOylation sites within the IKKβ protein sequence, supporting IKKβ as a potential SUMO-regulated target. Collectively, these analyses identify NF-κB signaling as a major SUMOylation-associated pathway and nominate IKKβ as a potential point of convergence between SUMOylation and inflammatory signaling.

### 3.14. CBO reprograms the immune transcriptional landscape during *K. pneumoniae* **infection**

An overview of host immune responses following infection and CBO treatment was obtained from the gene expression profile associated with SUMOylation, innate immune signaling, and inflammatory cytokines. Heatmap visualization demonstrated distinct transcriptional signatures across uninfected, infected, and CBO-treated macrophages **(Figure 5A)**. Infection with both classical *K. pneumoniae* (ATCC) and hypervirulent *K. pneumoniae* (hvKp) induced widespread activation of inflammatory genes while simultaneously altering the expression of key SUMO pathway components.

**Figure 5:**
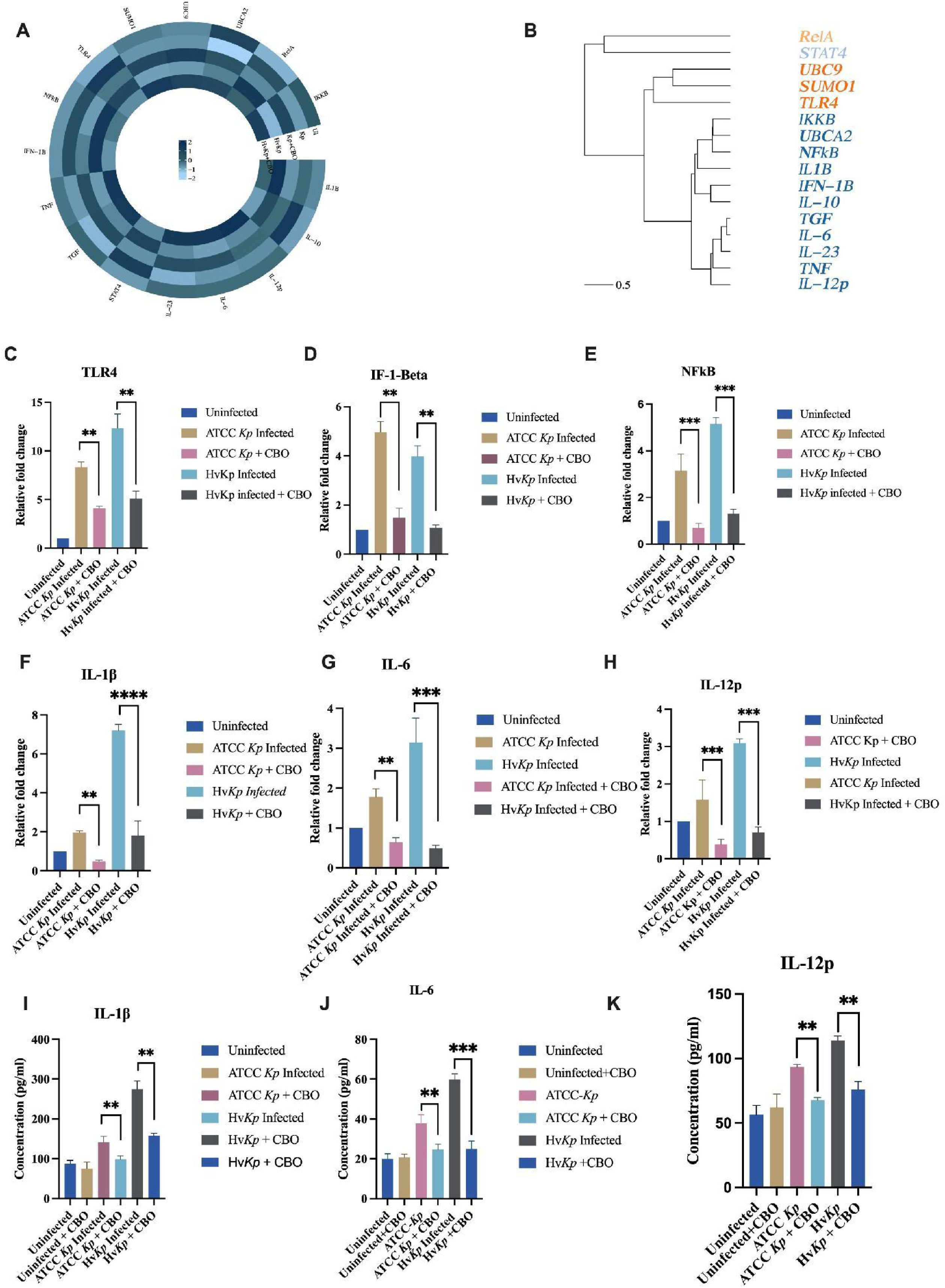
CBO suppresses infection-induced NF-κB activation and inflammatory responses during *Klebsiella pneumoniae* infection: (A) Heatmap showing expression patterns of SUMOylation-, NF-κB-, and cytokine-associated genes across experimental groups. (B) Hierarchical clustering analysis illustrates relationships among inflammatory and SUMOylation-related genes. (C–H) Relative mRNA expression of TLR4 (C), IFN-β (D), NFkB (E), IL-1β (F), IL-6 (G), and IL-12p (H) in THP-1 macrophages infected with classical *K. pneumoniae* (Kp) or hypervirulent *K. pneumoniae* (hvKp) following CBO treatment. Infection significantly upregulated inflammatory signaling genes, particularly in hvKp-infected cells, whereas CBO treatment reduced their expression toward basal levels. (I–K) ELISA quantification of IL-1β (I), IL-6 (J), and IL-12p (K) protein levels in culture supernatants. Consistent with transcriptional findings, infection elevated cytokine production, while CBO treatment significantly attenuated inflammatory cytokine secretion. Together, these data demonstrate that CBO mitigates infection-induced inflammatory responses and suppresses NF-κB-associated signaling during *K. pneumoniae* infection. Data are presented as mean ± SEM. Statistical significance is indicated as P < 0.01, P < 0.001, and P < 0.0001.

The hierarchical clustering further revealed coordinated regulation of functionally related genes **(Figure 5B)**. Genes involved in SUMOylation and upstream immune signalling, including SUMO1, UBC9, UBA2, TLR4, RelA, and IKBKB, clustered separately from downstream inflammatory cytokines, suggesting coordinated modulation of these signalling networks during infection. Notably, CBO treatment shifted the overall transcriptional profile away from the infection-associated state and towards that of uninfected macrophages, indicating a broad reprogramming of host immune responses rather than isolated changes in individual genes. These findings support the hypothesis that restoration of SUMOylation by CBO is accompanied by coordinated regulation of NF-κB-associated inflammatory pathways.

### 2.15. CBO suppresses infection-induced activation of TLR4/NF-κB-associated inflammatory genes

The global transcriptional changes of the selected genes involved in pathogen recognition, NF-κB signaling, and inflammatory responses were quantified by qPCR. Infection with both classical *K. pneumoniae* and hvKp significantly upregulated TLR4, IFN-β, NF-κB, IL-1β, IL-6, and IL-12p compared with uninfected controls **(Figures 5C–5H)**. Among the two strains, hvKp consistently elicited higher transcript levels, indicating stronger activation of innate immune signalling.

Treatment with CBO significantly attenuated the infection-induced expression of these genes in both infection models. Expression of TLR4, IFN-β, and NF-κB key showed an increase in expression level during infection, which was markedly reduced following CBO treatment, accompanied by decreased expression of downstream cytokine genes, including IL-1β, IL-6, and IL-12p. Although inflammatory gene expression was substantially reduced, transcript levels were not completely abolished, suggesting that CBO fine-tunes host immune activation rather than inducing broad immunosuppression. These findings are consistent with restoration of host SUMOylation and support a role for CBO in limiting excessive NF-κB-driven inflammatory responses during *K. pneumoniae* infection.

### 2.16. CBO restores cytokine homeostasis in infected THP-1-derived macrophages

To determine whether transcriptional changes translated into altered cytokine production, ELISA **(Figure 5I–5K)** quantified the secretion of inflammatory cytokines in culture supernatants. Consistent with the qPCR findings, infection with both classical *K. pneumoniae* and hvKp significantly increased the secretion of IL-1β, IL-6, and IL-12p, with hvKp producing a comparatively stronger inflammatory response. CBO treatment significantly reduced the elevated secretion of all three cytokines in infected macrophages. The reduction in cytokine production closely paralleled the corresponding decreases in mRNA expression, demonstrating that the transcriptional effects of CBO were reflected at the protein level. Importantly, cytokine production was moderated rather than completely suppressed, indicating that CBO restores immune homeostasis without compromising the capacity of macrophages to mount an inflammatory response. Collectively, these findings demonstrate that CBO reprograms both the transcriptional and functional inflammatory responses of macrophages during *K. pneumoniae* infection, supporting its role as a host-directed immunomodulatory agent.

### 2.16. Clove Bud Oil Modulates RelA and IKKβ Expression During *Klebsiella pneumoniae* Infection

To investigate the effect of CBO on NF-κB signaling during infection, the expression of RelA (p65) and IKKβ was evaluated at both protein and transcript levels in THP-1 macrophages infected with ATCC *K. pneumoniae* and hypervirulent *K. pneumoniae* (hvKp). Western blot analysis revealed increased RelA protein expression following infection with both ATCC and hypervirulent strains compared with uninfected controls **(Figure 6A, 6B)**. ATCC *K. pneumoniae* infection induced approximately a 3.2-fold increase in RelA expression, while hvKp infection produced a nearly 2.8-fold increase. Treatment with CBO significantly reduced RelA expression in both infection groups, lowering the levels to approximately 1.7-fold and 1.4-fold, respectively. In contrast, IKKβ protein expression was reduced during infection. Both ATCC *K. pneumoniae* and hvKp infection decreased IKKβ levels relative to uninfected cells **(Figure 6C, 6D)**. Following CBO treatment, IKKβ expression was restored and significantly increased compared with infected controls, reaching levels above those observed in uninfected cells, particularly in the hvKp-treated group. To further validate these findings, transcript levels of RelA and IKKβ were quantified by qPCR **(Figure 6E, 6F)**. RelA expression was markedly elevated following infection, showing approximately 11-fold and 18-fold increases in ATCC *K. pneumoniae* and hvKp groups, respectively. CBO treatment significantly reduced RelA transcript levels in both groups. Conversely, IKKβ expression was suppressed during infection but was restored following CBO treatment. ATCC *K. pneumoniae* and hvKp infection reduced IKKβ expression to nearly 0.5-fold of control levels, whereas CBO treatment increased expression to approximately 1.3-fold and 1.5-fold, respectively. Overall, both protein and gene expression analyses demonstrated that *K. pneumoniae* infection promotes activation of NF-κB signaling through increased RelA expression while suppressing IKKβ. Treatment with CBO reversed these infection-induced alterations by reducing RelA expression and restoring IKKβ levels, suggesting that CBO modulates NF-κB signaling and contributes to the restoration of immune homeostasis during *K. pneumoniae* infection.

**Figure 6:**
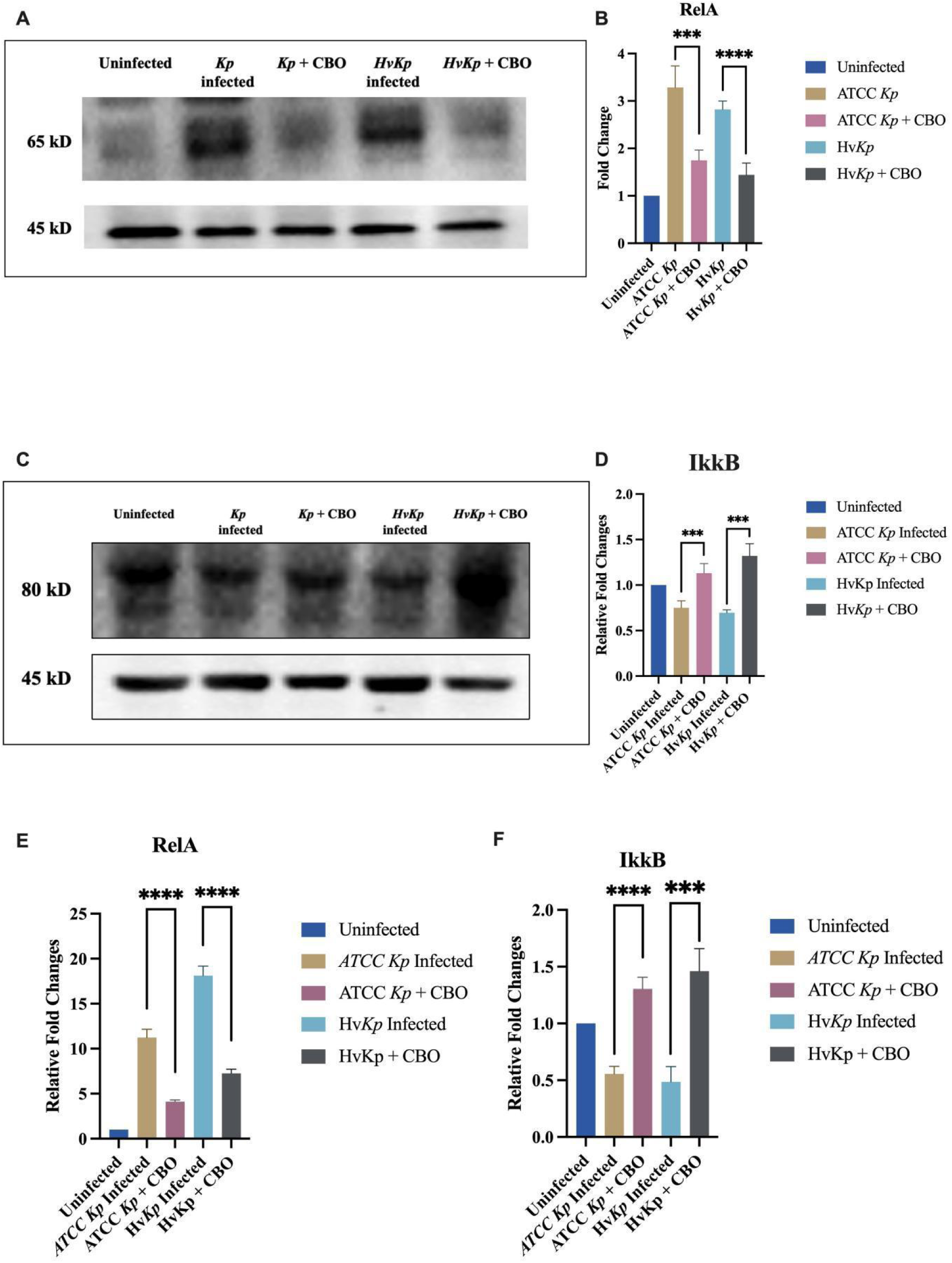
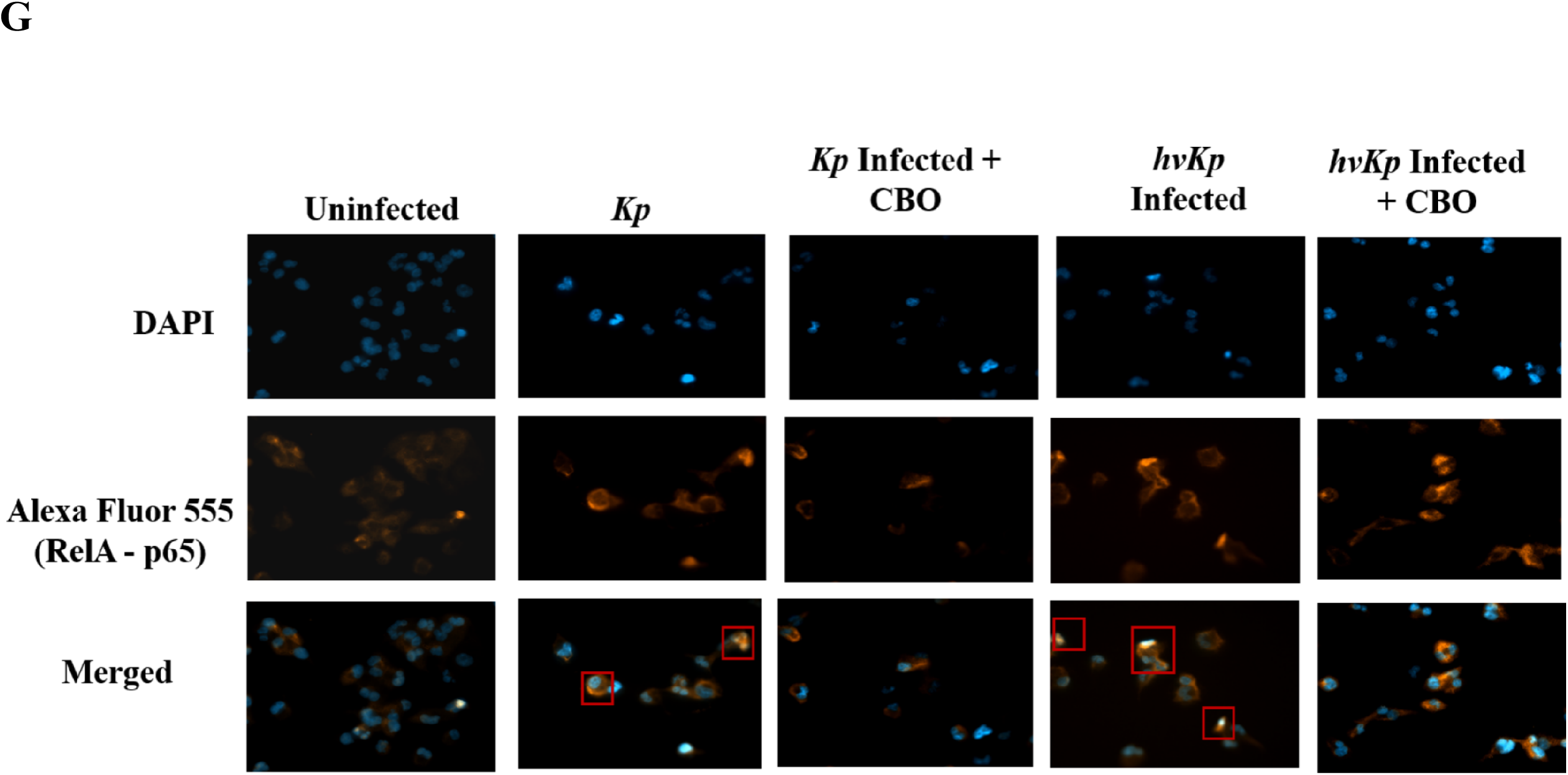
CBO modulates RelA and Ikkβ expression during *Klebsiella pneumoniae* infection: (A, B) Representative immunoblot and densitometric quantification of RelA (p65) expression in THP-1 macrophages infected with classical *K. pneumoniae* (Kp) or hypervirulent *K. pneumoniae* (hvKp). Infection increased RelA expression, whereas CBO treatment significantly reduced RelA levels in both infection models. (C, D) Representative immunoblot and densitometric quantification of Ikkβ expression. Infection reduced Ikkβ abundance compared with uninfected controls, while CBO treatment restored Ikkβ expression in both Kp- and hvKp-infected cells. (E, F) Relative mRNA expression of RelA (E) and Ikkβ (F) determined by qPCR. Consistent with protein-level observations, infection induced RelA expression and suppressed Ikkβ expression, whereas CBO treatment reversed these alterations. Together, these findings suggest that CBO attenuates infection-induced NF-κB activation while restoring IKKβ-associated signalling during *K. pneumoniae* infection. CBO reduces RelA (p65) nuclear localization during *Klebsiella pneumoniae* infection. Representative immunofluorescence images of THP-1-derived macrophages infected with classical *K. pneumoniae* (Kp) or hypervirulent *K. pneumoniae* (hvKp), with or without CBO treatment. Nuclei were stained with DAPI (blue), and RelA (p65) was detected using Alexa Fluor 555 (orange). Red boxes highlight representative cells showing nuclear RelA localization. Data are presented as mean ± SEM. Statistical significance is indicated as **P < 0.001 and *P < 0.0001.

### 2.17. CBO attenuates infection-induced nuclear localization of RelA during K. pneumoniae infection

To further investigate the effect of CBO on NF-κB signaling, the subcellular localization of RelA (p65) was examined by immunofluorescence microscopy. In uninfected macrophages, RelA staining was predominantly distributed outside the nucleus, indicating relatively low basal NF-κB activation. Infection with both classical *K. pneumoniae* and hvKp resulted in increased nuclear localization of RelA, as evidenced by enhanced overlap between Alexa Fluor 555-labeled RelA and DAPI-stained nuclei in the merged images. Representative cells displaying prominent nuclear RelA localization are highlighted by red boxes. In contrast, CBO treatment reduced the prominent infection-associated nuclear accumulation of RelA in both Kp- and hvKp-infected macrophages, with a greater proportion of the RelA signal appearing outside the nuclear compartment. These observations support the finding that CBO attenuates infection-induced RelA nuclear translocation and contributes to the normalization of NF-κB signaling. Together with the observed restoration of host SUMOylation and Ikkβ expression, these findings are consistent with a model in which CBO modulates the SUMOylation–NF-κB axis to restrain dysregulated inflammatory signaling during *K. pneumoniae* infection **(Figure 6G)**.

## 3. Materials and Methods

### 3.1. Reagents and Chemicals

Reagents and chemicals were purchased from HiMedia Laboratories, LLC, and Sigma Chemical Co. Fetal bovine serum (FBS) was obtained from Gibco Cell Culture Solutions (Thermo Fisher Scientific). Protease and phosphatase inhibitor cocktails were from Thermo Fisher Scientific. Antibodies against NFkB P65, RelA/NFkBphospho-IκBα, and anti-actin antibody were purchased from Cell Signalling Technologies (Danvers, MA, United States), and SUMO1 antibodies were purchased from Abcam (Cambridge, United Kingdom). The HRP-conjugated secondary antibodies were obtained from Cell Signalling Technologies (Danvers, MA, United States), and the enhanced chemifluorescence (ECF) reagent was obtained from Bio-Rad Laboratories. The polyvinylidene difluoride (PVDF) membranes were from Bio-Rad Laboratories (Hercules, CA, United States), Trizol® reagent was purchased from Invitrogen (Barcelona, Spain), and SYBR Green was obtained from Thermo Fisher Scientific. Primers were from Integrated DNA Technologies. CBO was obtained from Synthate Pvt. Ltd., and initial solubilization of CBO was performed in Dimethyl Sulfoxide (DMSO) (HIMEDIA, 67-68-5), then further solubilization in RPMI for the desired concentration of the oil. The remaining reagents were from Sigma Chemical Co. (Bayswater, VIC 3153, Australia).

### 3.2. Cell culture

THP1 cells were obtained from the ATCC American Type Culture Collection (TIB-71™-ATCC, USA). These cells were revived and maintained in a 75 cm^2^ culture flask in an incubator at 37±2°C and 5% CO₂. Cells were maintained in Dulbecco’s modified Eagle’s medium (RPMI, Gibco™, USA) supplemented with 10% fetal bovine serum (FBS, Gibco™, USA) and 1% penicillin and streptomycin antibiotic mixture (Invitrogen). Cell confluence was continuously monitored under an optical microscope until it reached approximately 75-80%. Monocytes were induced with 20 ng/ml phorbol-12-myristate-13-acetate (PMA; Sigma-Aldrich) for 24 h to differentiate into macrophages. This was confirmed by monitoring adherence and morphological changes using phase-contrast microscopy [13].

### 3.3. Isolation of Human PBMCs

Human PBMCs were isolated from peripheral blood collected in heparinized tubes from healthy volunteers. To enrich mononuclear cells, whole blood was diluted 1:1 with RPMI 1640 supplemented with 2% FBS and gently layered over an equal volume of Histopaque-1077. These samples were centrifuged at 600 g for 20 minutes at room temperature without a brake to allow clear separation of cell layers. A puffy, PBMC-enriched inferior layer was carefully separated and washed three times with 1X PBS. Recovered lymphocytes were counted and seeded for adherence. After 2 hours of incubation, non-adherent lymphocytes—including T cells, B cells, and NK cells—were removed, leaving behind the adherent monocytes, which were used for further experiments.

### 3.4. Bacterial strains and bacterial culture conditions

The study focuses on two *Klebsiella pneumoniae* strains: a reference strain, *K. pneumoniae* (ATCC 33495), from the American Type Culture Collection (ATCC, USA) and a clinical isolate, U10386 (hypervirulent), from Amrita Institute of Medical Sciences and Research Centre (AIMS, Kochi, India) [15]. The bacterial cultures were initiated by inoculating strains into Luria-Bertani (LB) broth (HiMedia, India) and incubating overnight at 37°C with constant agitation at 200 rpm. The overnight cultures were reinoculated at a 1:100 dilution into fresh LB broth to reach mid-log phase (OD600 = 0.4), as quantified using a biophotometer (Eppendorf, Germany). These cultures were used for all infection and viability assays [15].

### 3.5. Identification and characterization of clove bud oil

The global oil used in the study was obtained from Synthite Private Ltd., Kerala, India. According to the manufacturer’s specifications. The oil was obtained by steam distillation of the dried flower buds of *Syzygium aromaticum* (L.) Merr. & L.M. Perry (Family: Myrtaceae). The identity and chemical quality of the oil were verified using Fourier-transform infrared (FTIR) characterization previously performed in our laboratory and reported by H.J. et al. (2016). The FTIR spectrum exhibited the characteristic absorption band of CBO, confirming the chemical authenticity of the oil used in this study.

### 3.6. Bacterial Growth Kinetic Assay

The antibacterial activity of CBO against *Klebsiella pneumoniae* ATCC 33495 and hypervirulent *K. pneumoniae* (hvKp) strains was evaluated by monitoring bacterial growth kinetics. Overnight bacterial cultures were diluted in Luria–Bertani (LB) broth and dispensed into sterile 96-well microplates (100 µL per well). Various concentrations of CBO were added to the wells, while untreated cultures served as growth controls. Plates were incubated at 37°C in a Synergy H1 microplate reader (BioTek) with continuous orbital shaking. Bacterial growth was monitored by measuring optical density at 600 nm (OD600) at 30-min intervals over a 24-h period. Growth curves were generated by plotting OD600 values against time, and the effects of CBO on bacterial proliferation were compared with those of untreated controls [17].

### 3.6. Viability assessment

To determine CBO’s non-toxic concentration, its cytotoxicity was evaluated using the MTT assay. THP-1 monocytes (1 × 10⁵ cells/well) were seeded in 96-well plates and differentiated into macrophages using 20 ng/mL PMA for 24 h at 37°C in a 5% CO₂ incubator. Different concentrations of CBO ranging from 0.001% to 0.035% (V/V) were prepared and treated with cells for 24 and 48 h. Subsequently, 20 µL of MTT solution (5 mg/mL) was added to each well and incubated for 4 h. The resulting formazan crystals were dissolved in 100 μL DMSO, and absorbance was measured at 570 nm using a BioTek microplate reader.

### 3.7. Determination of minimum inhibitory concentration (MIC) of CBO on *K. Pneumoniae*

The CBO diluted with DMSO as the main stock and further diluted into a series of concentration gradients (0.01, 0.02, 0.03, 0.06, 0.125, 0.25, 0.5, 1.0, and 2%) was prepared to determine the MIC90 on *K. pneumoniae.* A 1:1 ratio of 0.4 OD bacteria (*K. pneumoniae):CBO* was added to a 96-well plate with sufficient shaking and further incubated at 37°C in a shaker for 24 h [18]. Similarly, we maintained a 100% oil, a 100% bacterial culture, and a bacterial control with DMSO, and the absorbance after 24 hours was measured at 600 nm using a Bio-Tek® microplate reader (Bio-Tek, Germany).

### 3.8. Intracellular reactive oxygen (ROS) species measurement

The THP-1-derived macrophages were co-infected with Kp and hvKp and treated with 0.015% of CBO for 24 h. Intracellular ROS levels were measured using DCFH-DA (Merck), a redox-sensitive dye. H_2_O_2,_ a positive control, was administered 30 minutes prior to the end of the treatment incubation, followed by 30 min incubation with DCFH-DA (10 μM) at 37°C for 45 min in the dark [19]. Fluorescence was measured using a microplate reader at excitation/emission of 485/535 nm.

### 3.9. Infection of THP-1-derived macrophages and CBO treatment

Differentiated macrophages were infected with *Klebsiella pneumoniae* strains with a multiplicity of infection (MOI) 1:15 for all the experiments performed [20]. Following a 30 min incubation for phagocytosis, cells were washed thrice with 1X Phosphate-Buffered Saline (PBS) and incubated in medium containing gentamicin (45 microgram/ml) for 45 min to remove the extracellular bacteria. CBO (0.015%) was added to appropriate wells, and cells were incubated for the preferred time as per each experiment. Uninfected macrophages with 0.5% of DMSO were always maintained as baseline controls in all the assays [21].

### 3.10. Phagocytosis and Intracellular Survival Assay

THP-1-derived macrophages were used to evaluate the effect of CBO on *Klebsiella pneumoniae* (Kp) and hypervirulent *K. pneumoniae* (hvKp) infection. For the pre-treatment group, cells were treated with 0.015% (v/v) CBO for 24 h before infection with Kp or hvKp (MOI 1:15). Following 30 min of phagocytosis, extracellular bacteria were removed by washing with PBS, and intracellular bacterial uptake was determined by lysing the cells with 0.06% SDS and enumerating colony-forming units (CFU). To assess intracellular bacterial clearance, infected cells were treated with gentamicin for 45 min to eliminate extracellular bacteria, then incubated in fresh medium containing 0.015% CBO for 24 h before CFU enumeration. For the post-treatment group, macrophages were infected for 30 min, washed, and subsequently treated with 0.015% CBO for 24 h prior to lysis and CFU determination.

### 3.11. Intracellular Survival Kinetics

THP-1-derived macrophages were pretreated with 0.015% (v/v) CBO for 24 h before infection. Following phagocytosis, extracellular bacteria were removed by 1X PBS washing, and intracellular bacterial uptake was determined by cell lysis (0.06% SDS) followed by CFU enumeration. To assess intracellular survival kinetics, cells were harvested immediately after treatment (0 h) and at 6-hour intervals through 48 h (0, 6, 12, 18, 24, 30, 36, 42, and 48 h). At each time point, cells were washed with 1X PBS and lysed with 0.06% SDS in 1X PBS for 5 minutes at room temperature to release intracellular bacteria. The lysates were serially diluted and plated onto Luria-Bertani (LB) agar, incubated overnight at 37 °C, and viable bacteria were enumerated as colony-forming units per milliliter (CFU/mL).

### 3.12. Blood Killing and Serum Killing

Human peripheral blood was collected from healthy volunteers into heparinized tubes. Whole blood was divided into active (37°C) and heat-inactivated (50°C, 30 min) aliquots, while serum was separated by centrifugation (2000 rpm, 15 min). Mid-log-phase *Klebsiella pneumoniae* (Kp) and hypervirulent *K. pneumoniae* (hvKp) cultures were inoculated into whole blood or serum, with or without CBO, and incubated at 37°C with gentle agitation. At 30-min intervals up to 2 h, aliquots were collected, host cells were lysed, and bacterial survival was determined by serial dilution and colony-forming unit (CFU) enumeration [6].

### 3.13. *C. elegans* Survival Assay to Assess the Anti-Infective Potential of Clove Bud Oil during *K. pneumoniae* infection

The anti-infective potential of CBO against *Klebsiella pneumoniae* infection was evaluated using a *Caenorhabditis elegans* survival assay. Wild-type N2 *C. elegans* were maintained on nematode growth medium (NGM) agar plates seeded with *Escherichia coli* OP50 as a food source under standard laboratory conditions. Age-synchronized L1 larvae were obtained and used for all experiments. For infection assays, groups of ten synchronized worms were transferred onto NGM plates seeded with either *K. pneumoniae* ATCC 33495 or a hypervirulent *K. pneumoniae* (HvKP) strain. To assess the protective effect of CBO, infected worms were exposed to CBO at the predetermined effective concentration. Control worms were maintained on *E. coli* OP50-seeded plates. Experimental groups included (i) E. *coli* OP50 control, (ii) *K. pneumoniae* infection, (iii) *K. pneumoniae* infection with CBO treatment, (iv) HvKP infection, and (v) HvKP infection with CBO treatment. Worm survival was monitored daily under a microscope. Worms that failed to respond to gentle mechanical stimulation were scored as dead and excluded from subsequent observations. Survival percentages were calculated for each group and compared to determine the protective effect of CBO against both classical and hypervirulent *K. pneumoniae* infections [22].

### 3.14. Gene expression studies of SUMO genes and other cytokines

Total RNA was isolated from THP-1 macrophages infected with Kp and hypervirulent Kp and treated with 0.015% CBO using the MACHEREY-NAGEL RNeasy Mini Kit® according to the manufacturer’s protocol. RNA purity and concentration were determined using a NanoDrop spectrophotometer. Samples were stored at -80°C until use. Complementary DNA (cDNA) was synthesized from 1 µg of RNA using the PrimeScript™ First Strand cDNA Synthesis Kit (Takara Bio, Japan) with incubation at 42 °C for 1 h followed by cooling to 4 °C. The synthesized cDNA was stored at −80 °C and used for quantitative PCR (qPCR) analysis. Primer specificity for SUMO isoforms and cytokine genes was validated by two-step RT-PCR. qPCR reactions were performed using SYBR® Green Master Mix (Applied Biosystems) on a QuantStudio™ Real-Time PCR System (Thermo Fisher Scientific) following the manufacturer’s instructions. Relative expression levels were calculated using the 2^(-ΔΔCt) method, with beta-actin as the internal control [23].

### 3.15. Western Blot Analysis

THP-1 macrophages were co-infected with *Klebsiella pneumoniae* (*Kp*) and hypervirulent *K. pneumoniae* (*HV-Kp*) at a multiplicity of infection (MOI) of 1:50 for 30 min to allow phagocytosis. Following infection, the macrophages were treated with 0.015% (v/v) CBO for 1 h, 3 h, and 5 h. Total cell lysates were prepared using 1× RIPA lysis buffer (Thermo Fisher Scientific, Cat. No. 89900) supplemented with protease inhibitor cocktail (Sigma-Aldrich, Cat. No. P8340). Protein concentration was determined using the bicinchoninic acid (BCA) protein assay kit (Thermo Fisher Scientific, Cat. No. 23225). Equal amounts of protein (40 µg) were resolved by SDS-PAGE on 12% gels and transferred onto nitrocellulose membranes (Bio-Rad, Cat. No. 1620112) using a semidry [23]. The membranes were blocked with 53% BSA in TBST (Tris-buffered saline with 0.1% Tween-20) for 1 h at room temperature and incubated overnight at 4°C with primary antibodies against SUMO1 (Abcam, Cat. No. ab32058; 1:1000) and UBC9 (Abcam, Cat. No. ab3742; 1:1000). After washing, membranes were incubated with HRP-conjugated anti-rabbit IgG secondary antibody (Cell Signaling Technology, Cat. No. 7074; 1:10,000) for 1 h at room temperature. Protein bands were visualized using enhanced chemiluminescence (ECL) reagent (Bio-Rad, Cat. No. 1705061) and imaged on a Bio-Rad Gel Doc imaging system. The blots were stripped and reprobed with anti-β-actin antibody (Sigma-Aldrich, Cat. No. A5316; 1:5000) as the loading control. For NF-κB pathway analysis, a similar experimental setup was followed. THP-1 macrophages were co-infected with Kp and HV-Kp and treated with 0.015% CBO for 1 h, 3 h, 5 h, and 24 h. Cell lysates were prepared as described above, and 30 µg of total protein was separated by SDS-PAGE and transferred onto nitrocellulose membranes. The membranes were probed with antibodies against RelA (p65) (Cell Signaling Technology, Cat. No. 8242; 1:1000) and IKKβ (Cell Signaling Technology, Cat. No. 8943; 1:1000), followed by HRP-conjugated secondary antibody. Immunoreactive bands were detected using ECL and visualized using the Bio-Rad Gel Doc imaging system. The same blots were stripped and reprobed with β-actin as an internal loading control [24].

### 3.16. Identification of SUMOylation-associated inflammatory signaling pathways

To investigate signaling pathways associated with the SUMOylation machinery, proteins involved in SUMO conjugation and deconjugation, including SUMO1, SUMO2, UBA2, UBE2I, PIAS family proteins, and SENPs, were selected for analysis. Protein interaction networks were generated using the STRING database (v12.0), and network visualization was performed using Cytoscape (v3.10) [25].

### 3.16.1. Functional Enrichment Analysis

Functional enrichment analysis was performed using the STRING enrichment module. Gene Ontology (GO) biological process and Kyoto Encyclopedia of Genes and Genomes (KEGG) pathway analyses were conducted to identify biological pathways associated with the selected SUMOylation-related proteins. Significantly enriched pathways were identified based on false discovery rate (FDR)-adjusted p-values.

### 3.16.2. Pathway Interaction Analysis

To explore potential signaling pathways associated with SUMOylation, pathway interaction analyses were performed using Pathway Commons and the Reactome Pathway Browser. Curated molecular interactions involving SUMOylation-associated proteins and their interacting partners were examined to identify potential signaling networks associated with SUMO-dependent regulation.

### 3.16.3. Protein–Protein Interaction Network Construction

Proteins identified from pathway enrichment and interaction analyses were further subjected to protein–protein interaction (PPI) network analysis using STRING. The resulting interaction networks were visualized and analyzed in Cytoscape to examine network connectivity and identify key interaction nodes linking SUMOylation-associated proteins with inflammatory signaling mediators.

### 3.16.4. Prediction of SUMOylation Sites in IKKβ

Based on the pathway and network analyses, IKKβ (IKBKB) was selected for further investigation as a potential SUMOylation target. The amino acid sequence of human IKKβ (UniProt ID: O14920) was retrieved from the UniProt database and analyzed using the GPS-SUMO prediction server. Predicted SUMOylation sites were identified using the default prediction parameters and mapped onto the IKKβ protein sequence for further analysis.

### 3.17. Cytokine Quantification by Enzyme-Linked Immunosorbent Assay (ELISA)

The cytokine secretion was analyzed in THP-1 differentiated macrophages across five experimental groups: uninfected control, cells infected with *Klebsiella pneumoniae* (*Kp*), cells infected with hypervirulent *K. pneumoniae* (*HV-Kp*), and infected cells treated with CBO (*Kp* + CBO and *HV-Kp* + CBO). Macrophages were co-infected at a multiplicity of infection (MOI) of 50 for 30 min to allow for phagocytosis. Following infection, extracellular bacteria were removed, and cells were incubated in media containing 0.015% (v/v) CBO for 24 h. Cell-free culture supernatants were subsequently collected and analyzed for the release of IL-1B, IL-6, IL-12, IL-23, TNF-alpha, TGF-B, and IL-10. Quantification was performed using commercially available Sandwich ELISA kits (Origin Technologies, India) according to the manufacturer’s instructions. Absorbance was measured at 450 nm using a microplate reader. Cytokine concentrations were determined by interpolation from a standard curve generated using standards provided in the kits [25].

### 3.18. Immunofluorescence analysis of RelA (p65) nuclear localization

THP-1 monocytes were seeded in imaging plates and differentiated into macrophages using 20 ng/mL phorbol 12-myristate 13-acetate (PMA) for 24 h. Differentiated macrophages were infected with either Klebsiella pneumoniae ATCC 33495 (Kp) or hypervirulent K. pneumoniae (hvKp) at the selected multiplicity of infection for 30 min to allow bacterial uptake. Following infection, extracellular bacteria were removed by washing with 1× PBS, and the cells were treated with 0.015% (v/v) CBO under the indicated experimental conditions. Following treatment, cells were washed with 1× PBS and fixed with 4% paraformaldehyde. The cells were subsequently permeabilized with 0.1% Triton X-100 and blocked with 5% goat serum to minimize nonspecific antibody binding. Cells were incubated with a primary antibody against RelA (p65) at a dilution of 1:400, followed by incubation with an Alexa Fluor 555-conjugated secondary antibody at a dilution of 1:500. Nuclei were counterstained with DAPI. Fluorescence images were acquired using an ApoTome fluorescence imaging system under identical acquisition settings across all experimental groups. The experimental groups included uninfected control, Kp-infected, Kp-infected + CBO, hvKp-infected, and hvKp-infected + CBO macrophages. Nuclear localization of RelA was assessed based on the spatial overlap between the Alexa Fluor 555 RelA signal and DAPI-stained nuclei in merged fluorescence images.

### 3.18. Statistical analysis

All data are presented as the mean ± standard error of the mean (SEM). Statistical analyses were performed using GraphPad Prism software. The significance of differences between multiple experimental groups was determined using a one-way analysis of variance (one-way ANOVA)/two-way analysis of variance (two-way ANOVA). A post-hoc test was subsequently used to identify significant differences between specific pairs of groups. In all analyses, a P-value of less than 0.05 was considered statistically significant.

## 4. Discussion

Post-translational modifications (PTMs) are fundamental regulators of cellular physiology and innate immunity during microbial infection [26]. Among these, SUMOylation is a dynamic and reversible modification that controls protein stability, subcellular localization, transcriptional activity, and protein–protein interactions. Intracellular pathogens manipulate this machinery through distinct mechanisms: *Listeria monocytogenes* induces degradation of the SUMO-conjugating enzyme UBC9, suppressing global host SUMOylation [27], whereas *Salmonella enterica* exploits SUMOylation of its effector SifA to maintain the Salmonella-containing vacuole [28]. Effectors from *Anaplasma phagocytophilum* [29] and *Ehrlichia chaffeensis* [30] similarly undergo host-mediated SUMOylation to support intracellular survival. *Klebsiella pneumoniae* also perturbs host immunity to promote persistence [31, 32], with previous studies implicating IFN-β-dependent induction of let-7 microRNAs, reduced SUMO1 expression, and SENP2 activation in this process [10]. However, the mechanism by which restoration of host SUMOylation can restore antibacterial immunity during *K. pneumoniae* infection remains poorly understood.

The CBO concentration used in functional assays was well tolerated by THP-1 macrophages and remained below the minimum inhibitory concentration for both classical and hypervirulent strains, indicating that its antibacterial effects were unlikely to arise from direct growth inhibition. CBO increased bacterial uptake during early infection and subsequently reduced intracellular survival, suggesting enhancement of both phagocytosis and bacterial killing. This activity is particularly relevant in the case of hypervirulent *K. pneumoniae*, as the presence of its prominent capsule and additional immune-evasion mechanisms impede phagocytic clearance. Treatment also restored intracellular ROS production, providing a plausible mechanism for the observed reduction in bacterial burden, as the oxidative burst constitutes a principal antimicrobial function of professional phagocytes.

A central finding was the coordinated suppression of the host SUMO-conjugation machinery during infection. Reduced expression of SUMO1, UBA2, and UBC9 was accompanied by diminished global SUMO conjugation, suggesting that *K. pneumoniae* interferes with the pathway at multiple regulatory levels. CBO reversed these alterations, restoring key SUMO-pathway components and cellular conjugation capacity. Given the involvement of SUMOylation in transcriptional regulation, inflammatory signalling, vesicular trafficking, and antimicrobial defense, recovery of this machinery may influence several immune processes simultaneously. This mechanism differs from the exploitation of an individual SUMOylated bacterial effector, such as *Salmonella* SifA [28], and instead suggests a broader disruption of host post-translational regulation by *K. pneumoniae*. SUMOylation shapes immune responses through several regulatory mechanisms, with the NF-κB pathway constituting a major regulatory axis [33]. SUMOylation modulates the stability, activity, and subcellular localization of key regulators, including IKKβ and RelA, thereby controlling IκB turnover, RelA nuclear translocation, and inflammatory gene expression [34]. However, suppression of SUMOylation during *K. pneumoniae* infection was associated with enhanced IKKβ, resulting in sustained RelA activation and enhanced pro-inflammatory signalling [35, 36]. CBO restored SUMO-pathway components and IKKβ abundance while constraining RelA activation and downstream inflammatory responses. These observations support functional crosstalk between SUMOylation and NF-κB signalling, although the relevant SUMO-modified substrates and direct molecular interactions remain to be defined.

CBO selectively attenuated the infection-induced elevation of IL-1β, IL-6, IL-12p40, and IFN-β [37]. While enhancing bacterial clearance. This response suggests restoration of immune homeostasis, permitting effective antimicrobial activity without sustaining potentially damaging inflammation. The concurrent improvement in phagocytosis, ROS production, intracellular killing, and complement-dependent elimination further indicates that CBO reinforces multiple arms of macrophage defense. Nevertheless, direct dependence of these effects on SUMOylation requires confirmation through genetic or pharmacological disruption of individual SUMO-pathway components.

Together, these findings support a model in which *K. pneumoniae* suppresses host SUMOylation, potentially through an IFN-β-associated regulatory axis, thereby perturbing IKKβ–RelA signalling and inflammatory homeostasis. CBO reverses this molecular phenotype by restoring SUMO-conjugation capacity, recalibrating NF-κB-associated responses, and strengthening oxidative and complement-mediated antibacterial activity, ultimately restricting intracellular survival of classical and hypervirulent *K. pneumoniae* (Fig. 7). This study therefore links restoration of host SUMOylation to the immunomodulatory activity of CBO and identifies SUMO-dependent signalling as a potential target for host-directed intervention. Identification of the relevant SUMO-modified substrates, together with temporal and loss-of-function studies, will be essential to establish causality and determine the translational potential of this strategy.

**Figure 7:**
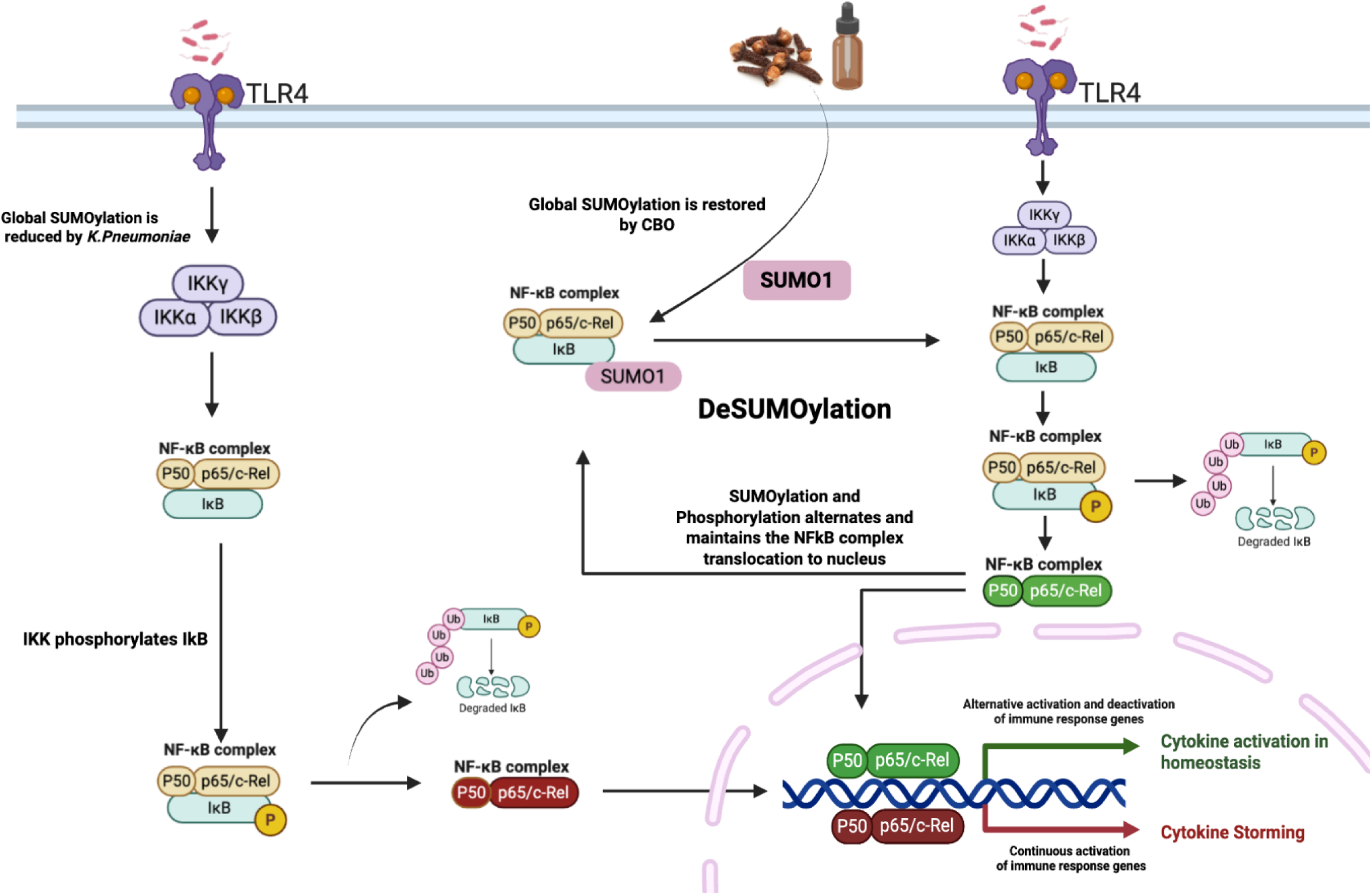
Proposed model illustrating SUMOylation-mediated regulation of NF-κB signaling during *Klebsiella pneumoniae* infection and its restoration by Clove Bud Oil (CBO): Under physiological conditions, balanced SUMOylation maintains regulated IKKβ phosphorylation, controlled NF-κB activation, and immune homeostasis. During *K. pneumoniae* infection, activation of the IFN–IFNR–let-7 axis suppresses the host SUMOylation machinery, resulting in reduced SUMOylation, persistent Ikkβ phosphorylation, enhanced RelA (p65) nuclear accumulation, and excessive NF-κB activation. These alterations promote dysregulated inflammatory responses characterized by increased expression of pro-inflammatory cytokines and impaired immune homeostasis. CBO treatment restores the expression of SUMOylation-associated components, enhances SUMO conjugation, and re-establishes balanced regulation of Iκκβ and NF-κB signalling. Restoration of SUMOylation limits aberrant RelA activation normalizes cytokine expression, promotes immune homeostasis, and enhances bacterial clearance. This model summarizes the proposed mechanism by which CBO modulates host SUMOylation–NF-κB signaling during both classical and hypervirulent *K. pneumoniae* infection.

## Ethics statement

Blood was collected via venipuncture from healthy volunteers under written informed consent approved by the Ethics Committee of Amrita School of Medicine (ECASM-AIMS-2023-149), Amrita Vishwa Vidyapeetham. All experiments were performed in accordance with relevant guidelines and regulations of Amrita School of Biotechnology, Amrita Vishwa Vidyapeetham, India.

## Acknowledgement

The authors gratefully acknowledge Sri Mata Amritanandamayi Devi, Chancellor of Amrita Vishwa Vidyapeetham, for her constant support, guidance, and funding for the research. The author also acknowledges the doctoral fellowship from UGC—the Savitribai Jyotirao Phule Fellowship for a Single Girl Child.

